# Possible Mechanism of Porosome-Mediated Exosome Release

**DOI:** 10.1101/2022.03.06.483164

**Authors:** Won Jin Cho, Brent Formosa, Mzia G. Zhvania, Douglas J. Taatjes, Ionita Ghiran, Bhanu P. Jena

## Abstract

Supramolecular cup-shaped lipoprotein structures called *porosomes* embedded in the cell plasma membrane mediate fractional release of intra-vesicular contents from cells during secretion. The presence of porosomes, have been documented in many cell types including neurons, acinar cells of the exocrine pancreas, growth hormone secreting cells of the pituitary, and mouse insulin-secreting beta cell insulinomas called Min6. Electron micrographs of the nerve terminal in rat brain and of Min6 cells, suggest that besides the docking, fusion and content release of neurotransmitter containing synaptic vesicles and insulin containing granules, multi-vesicular bodies and exosome-like vesicles are released at the porosome complex. To test this hypothesis, the porosome-associated calcium transporting ATPase 1 [SPCA1] encoded by the ATP2C1 gene, was knocked-out in Min6 cells using CRISPR, to study glucose-stimulated insulin and exosome release. In agreement with electron micrographs, results from the study demonstrate a loss of both glucose-stimulated insulin and exosome release in the ATP2C1 knockout Min6 cells. These results further confirm the role of the porosome complex in insulin secretion and establishes a new paradigm in porosome-mediated exosome release in beta cells of the endocrine pancreas.

## INTRODUCTION

Nearly 38 trillion cells composing the human body communicate with each other via secreted biomolecules. In 1996, plasma membrane-associated 100-120 nm cup-shaped lipoprotein structures called *porosomes* were discovered in the exocrine pancreas and subsequently in the endocrine cells and neurons, that enable the secretion of biomolecules from cells (1–13). Porosomes in neurons are approximately 15 nm cup-shaped lipoprotein structures with a central plug, and are composed of nearly 30 proteins (7,9,10). Porosomes mediate the kiss-and-run mechanism of fractional release of intravesicular contents during cell secretion (3–10). In porosome-mediated secretion, secretory vesicles temporarily dock and fuse at the porosome base in the cell plasma membrane as opposed to complete collapse of the vesicle membrane with the cell plasma membrane to empty intravesicular contents during secretion. Glucose stimulated release of insulin stored in secretory vesicles in β-cells occur either by complete collapse of the vesicle membrane at the cell plasma membrane or the transient docking and fusion of secretory vesicles at the base of porosomes. In agreement therefore, partially empty secretory vesicles accumulate in cells following secretion, as demonstrated in electron micrographs (11). In earlier studies, isolated porosomes from the exocrine pancreas and neurons, have been structurally and functionally reconstituted into artificial lipid membranes and live insulin-secreting mouse insulinoma cell line, Min6 (4,7,11,12). In the past two decades, a large body of information provide conclusive evidence of secretory defects resulting from mutation in genes expressing various porosome proteins (13). Among these proteins, is the calcium transporting ATPase 1 [SPCA1] encoded by the ATP2C1 gene. Glucose stimulated release of insulin stored in secretory vesicles in β-cells occur either by complete collapse of the vesicle membrane at the cell plasma membrane or the transient docking and fusion of secretory vesicles at the base of porosomes. To introduce the reader on the structure of the porosome complex in the exocrine and endocrine pancreas and neurons (2,4,7), Figure 1 is presented, with secretory vesicles and the porosomes depicted in pseudo color for clarity.

**Figure 1.**
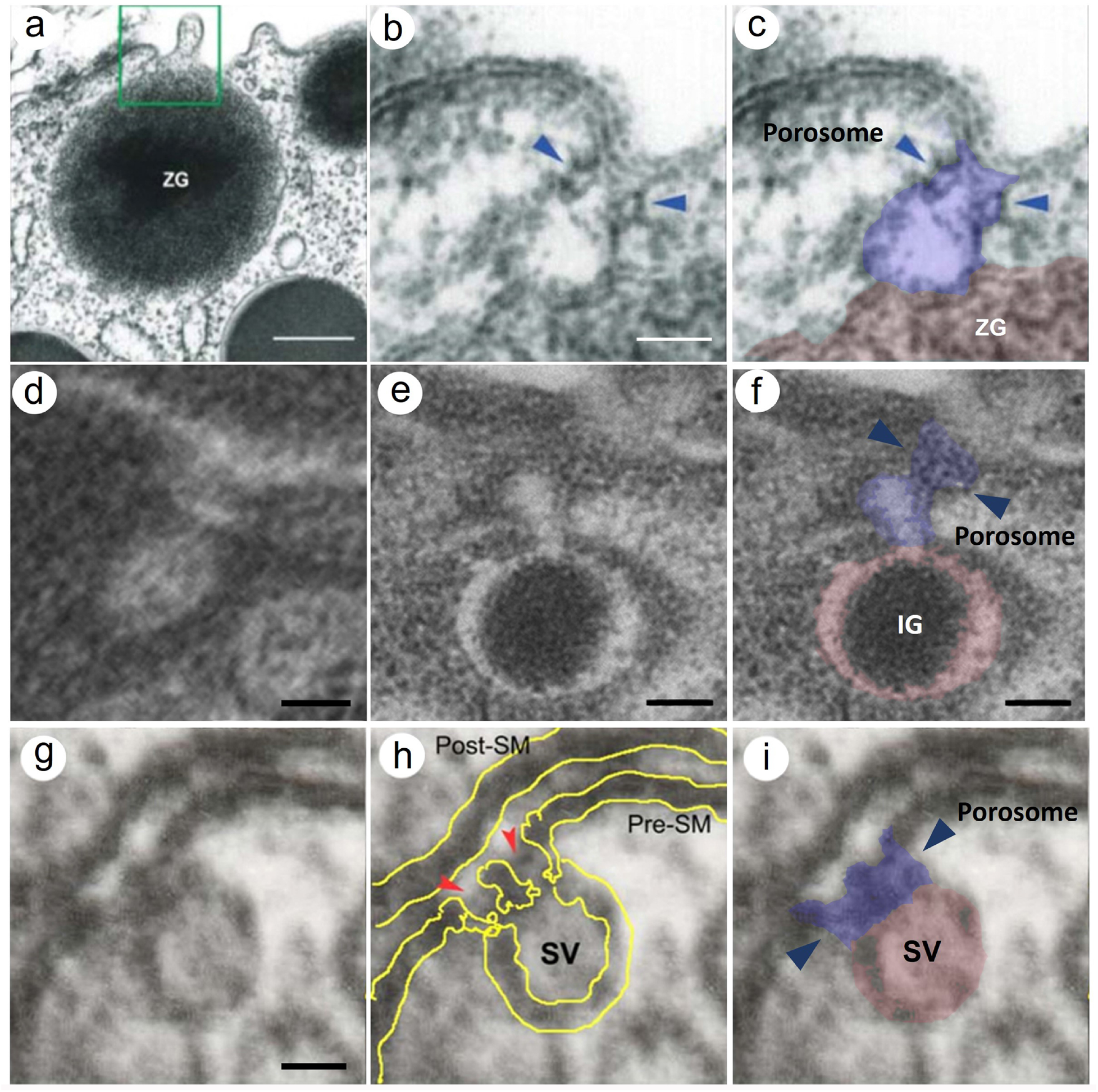
Electron micrographs of cup-shaped porosome complexes (purple) with docked secretory vesicles (pink) at the cell plasma membrane in the exocrine pancreas **(a-c)**, insulin secreting beta cells of the endocrine pancreas **(d-f)** and at the nerve terminal in neurons **(g-i)**. (**a**) Electron micrograph of the apical region of an exocrine pancreatic acinar cell, showing within the green boxed area, a membrane bound secretory vesicle called zymogen granule (ZG) docked at the base of a 100 nm cup-shaped porosome complex at the cell plasma membrane (*Scale bar = 400 nm*). Note at greater magnification **(b, c)**, (*Scale bar = 50 nm*) the plasma membrane bilayer, the bilayered cup-shaped porosome and the bilayer membrane of ZG are observed. Also note the ring complex at the neck of the porosome (blue arrow heads), proposed to enable the regulated opening and closing of the porosome complex, act via actin and myosin motor proteins. Similarly, cup-shaped porosome complexes are present in the insulin-secreting cells of the endocrine pancreas **(d-f)** (*Scale bar = 100 nm*). Note a 250 nm insulin secreting granule (IG) docked, fused and in the process of releasing insulin via the porosome in Min6 cells. **(g-i)** Cup-shaped 15 nm neuronal porosome complexes at the nerve terminal. Note the central plug within the neuronal porosome complex. (*Scale bar = 15 nm*). Parts of the images are adopted from our earlier studies (2,4,7), and presented for clarity.

In addition to secretion of neurotransmitters, digestive enzymes or hormones, cells also communicate with each other via secreted membrane-bounded nanostructured extracellular vesicles called *exosomes*, first discovered and reported in 1983 as a selective externalization mechanism of the transferrin receptor in sheep reticulocytes (14). Electron micrographic evidence for such externalization of the transferrin receptor in vesicular form from sheep reticulocytes, was demonstrated in 1985 (15). In the past 35 years, great progress has been made in our understanding of the biology, function, and biomedical application of exosomes (16–18). Exosomes vesicles are packaged with proteins, DNA, and RNA, that are destined for specific target cells in the body. Such intercellular communication via exosomes has also been implicated in various pathologies such as cancers, neurological disorders and inflammatory disease (16–18). While extracellular vesicle cargo include plasma membrane and endosomal proteins, they may also contain materials from various intracellular compartments such as the mitochondria. Recent studies report the presence of mitochondrial DNA within extracellular vesicles (19, 20). Although, multi-vesicular bodies may fuse at the cell plasma membrane to release their cargo, the molecular mechanism of exosome release and or their cargo from various cell types, remains unclear.

The current study conducted in rat brain tissue and Min6 cells using electron microscopy (EM), suggests the possible release of exosomes via the porosome complex. Syntaxin and the ATP2C1 gene product are among the nearly 30 porosome-associated proteins. To further test whether porosome are indeed involved in the release of exosomes in Min6 cells, glucose stimulated insulin secretion assays were performed in control wild type (SCRM) and in ATP2C1 gene knocked-out Min6 cells using CRISPR. Results from the study confirms the release of exosomes via the porosome complex as observed in EM micrographs, demonstrating a loss of both glucose-stimulated insulin secretion and exosome release in the ATP2C1 knockout Min6 cells. These results further establishes a new paradigm in porosome function in neurons and beta cells of the endocrine pancreas.

## MATERIALS AND METHODS

### Rat brain tissue sections

As previously published (7), Wistar rats, weighing approximately 115g were used in the study. Animals were housed in a controlled environment (temperature 20-22 °C, 60% humidity, 12h light/dark cycle) with free access to food and water. Following euthanasia using pentobarbital (100 mg/kg), animals underwent trans-cardiac perfusion with phosphate buffer saline pH 7.4 containing 2.5% glutaraldehyde, prior to processing for electron microscopy (7).

### Min6 cell culture

Min6 mouse insulinoma cells were cultured according to published procedure (2) in high-glucose (25 mM) Dulbecco’s Modified Eagle Medium (DMEM) (Invitrogen) supplemented with 10% fetal calf serum, 50 μM β-mercaptoethanol and antibotics (Penicillin and Streptomycin). Porosome isolations and electron microscopy were performed using Min6 cells grown to confluence in 100 x 13 mm sterile plastic petri dishes. Immunofluorescence microscopy was performed on Min6 cells grown to 75% confluence in 35 mm petri dishes with glass bottom coverslips (MatTek, Ashland, MA).

### Min6 cell ATP2C1 knockout using CRISPR

ATP2C1 knock out Min6 cells were generated by genome editing using a CRISPR/Cas9 system. In brief, pLentiCRISPR v2 plasmids that contained predesigned guide RNA targeting mouse ATP2C1 (KO, 5′-TGATGCCGTCAGTATCACTG-3′) and scrambled control guide RNA (SCRM, 5’- AAACCAAAGAGCCGAAGAAC-3’) were obtained from GenScript (Piscataway, NJ). These plasmids were then transfected into Min6 using Lipofcetamin 3000 (Invitrogen). Cells were selected with puromycin (0.15ug/ml) for 10 days until resistant populations emerged. After clonal expansion of each cell, the knockout of ATP2C1 was confirmed via immunoblot.

### PCR

The aliquots of each condition were collected, followed by PCR analysis using 2X DreamTaq Green PCR Master Mix (2X) (Thermo Fisher Scientific) with random hexamer (Invitrogen) as PCR primers.

### Statistical analysis

All experiments were repeated at least three times, and all data are presented as the mean ± SD. Statistical analysis was performed using student unpaired two-tailed t-test whose significance was determined according to p values (p<0.01(**), p< 0.05(*)), no significance to n.s. between studied groups, respectively.

### Electron microscopy (EM)

#### Rat brain EM

Transmission electron microscopy of rat brain was performed as described in a previously published procedure (7). Briefly, rat brain was perfused with normal saline solution, followed by phosphate buffer (pH 7.4) containing 2.5% glutaraldehyde. After perfusion, the brain was carefully removed and diced into 1 mm^3^ pieces. The pieces of brain tissue were post-fixed in phosphate buffer containing 1.5% osmium tetroxide, dehydrated in graded ethanol and acetone, and embedded in araldite. Tissue blocks were appropriately trimmed and the 40-50 nm sections obtained were stained with lead citrate and examined under a JEOL JEM-100C transmission electron microscope.

#### Min6 Cell EM

Transmission electron microscopy of Min6 cells was performed as described in a previously published procedure (8,9). Briefly, cells were fixed in 2% glutaraldehyde/2% paraformaldehyde in ice-cold phosphate buffered saline (PBS) for 24 h, washed with buffer, embedded in 2% SeaPrep agarose, followed by post-fixation for 1 h at 4 °C using 1% OsO_4_ in 0.1 M cacodylate buffer. The sample was then dehydrated in a graded series of ethanol, through propylene oxide, and infiltrated and embedded in Spurr’s resin. Ultrathin sections were cut with a diamond knife, retrieved onto 200 mesh nickel thin-bar grids, and contrasted with alcoholic uranyl acetate and lead citrate. Grids were viewed with a JEOL 1400 transmission electron microscope (JEOL USA, Inc., Peabody, MA) operating at 60 or 80 kV, and digital images were acquired with an AMT-XR611 11 megapixel CCD camera (Advanced Microscopy Techniques, Danvers, MA).

### Pseudo coloring of electron micrographs

Transmission electron micrographs of Min6 cells were pseudo colored for clarity using the published Basic Photoshop for Electron Microscopy approach [http://www.nuance.northwestern.edu/]. The main purpose of this approach is to yield additional valuable information for clarity and structural details that could easily be overlooked. The original EM images are provided alongside the pseudo-colored images, for further authenticity and clarity.

### Immunofluorescence microscopy

To determine the presence and distribution of insulin and the porosome-associated proteins Syntaxin and ATP2C1 in control (SCRM) and ATP2C1-KO Min6 cells, immunofluorescence studies were performed according to published procedures (8,14). To determine the position of the cell nucleus, cells were exposed to DAPI nuclear stain (Molecular Probes, Life Technologies, Carlsbad, CA). Immunofluorescent images were acquired using an immunofluorescence FSX100 Olympus microscope through a 100x objective lens (numerical aperture = 1.40) with illumination at 405 nm, 488 nm, or 647 nm. The co-association of Syntaxin 1 and ATP2C1 and their cellular distribution was determined by merging the fluorescent images.

### Fluorescent Immuno-colocalization Analysis

Image analysis of colocalization was performed using JACop tool of Plugins in the ImageJ (v1.53p). The comparative degree of colocalization for the SCRM and ATP2C1 KO was calculated as mean Pearson’s coefficients on the red and green channels.

### Densitometric Analysis of Immunoblots

Densitometry analysis of immunoblots was performed using the ImageJ (v1.53p) program. Values were adjusted to the proper experimental conditions then the values were displayed under each panel as a fold change.

### Glucose-stimulated insulin secretion from Min6 cells

MIN6 cells grown to 75% confluency in 100 × 13 mm sterile plastic petri dishes and glucose-stimulated insulin release was estimated. All secretion assays were performed at room temperature (25 °C). Cells were washed three times using 5ml/wash of PBS, pH 7.4, and incubated in 35 mM glucose-PBS. Since 30 mM glucose demonstrated no significant change in insulin secretion over control, and 35 mM glucose did (data not shown), 35 mM glucose was used in all assays carried out in this study. Two hundred microliter aliquots were removed at times 0, 10, and 30 min following 35 mM glucose incubation. The aliquots were centrifuged at 4,000 x*g* to remove any cells that may have been aspirated, and 160 μl of the supernatant was mixed with 40 μl of 5x Laemmli reducing sample preparation buffer (13), boiled for 2 min, and resolved using SDS-PAGE followed by Western blot analysis utilizing an insulin-specific antibody. Following completion of the secretion assays, cells were solubilized in equal volumes of PBS, their protein concentration determined. To compare total insulin in the control and reconstituted cells, equal volume of the cell lysate in Laemmli reducing sample preparation buffer (13) was immunoblotted using insulin-specific antibodies. Five micrograms of the cell lysate was also used in SDS-PAGE and Western blot analysis to determine the immunoreactive presence of various porosome-associated proteins. Percent insulin release were measured from the optical densities of insulin Western blots of the secreted and whole cell lysates.

### Western blot analysis

Isolated Min6 homogenates from both control (SCRM) and ATP2C1-KO and the glucose-stimulated insulin release medium in Laemmli buffer were resolved in a 12.5% SDS-PAGE, followed by electrotransfer to 0.2 mm nitrocellulose membrane. The membrane was incubated for 1h at room temperature in blocking buffer (5% nonfat milk in PBS pH 7.4 containing 0.1% Triton X-100 and 0.02% NaN_3_) and immunoblotted for 2 h at room temperature with antibodies raised against insulin (Santa Cruz Biotechnology Inc, Santa Cruz, CA), the porosome associated protein ATP2C1 and actin (Santa Cruz Biotechnology Inc, Santa Cruz, CA), and the exosome marker CD63 (Santa Cruz Biotechnology Inc, Santa Cruz, CA). All antibodies were used at a final concentration of 0.2μg/ml in blocking buffer. The immunoblotted nitrocellulose sheets were washed in PBS (pH 7.4) containing 0.1% Tween, prior to incubation for 1h at room temperature in horseradish peroxidase-conjugated secondary antibodies at a dilution of 1:5,000 in blocking buffer. The immunoblots were washed in PBS containing 0.1% Tween and processed for enhanced chemiluminescence and exposure to X-Omat-AR film. The exposed films were then developed and photographed.

## RESULTS AND DISCUSSION

In the current study, close examination of electron micrographs of rat brain tissue and insulin secreting Min6 cells, reveal the possible release of exosome-like vesicles measuring 15-20 nm in diameter and or the secretion of their intra-vesicular contents, via the porosome complex. To test the hypothesis that porosome are involved in exosome secretion, glucose stimulated insulin and exosome secretion from Min6 cells were performed in wild type SCRM control and in ATP2C1 gene knocked-out Min6 cells using CRISPER. Results from the study confirms the release of both insulin and exosomes via the porosome complex in Min6 cells.

### Electron micrographs of rat brain tissue suggests cargo release from exosome-like vesicles via fusion with synaptic vesicles docked at the porosome complex

Electron micrograph of rat brain tissue [Figure 2] shows the presence of 15-20 nm exosome-like vesicles free or associated with 30–40 nm synaptic vesicles. Some of the exosome-like vesicles appear to be generated from the mitochondria present within the synaptosome. Exosome-like vesicles are also seen associated with synaptic vesicles docked at the neuronal porosome complex, suggesting fusion and content release together with neurotransmitters. Multiple exosome-like vesicles are sometimes found associated with a single synaptic vesicle. Similar to the clear synaptic vesicles, the exosome-like vesicles appear clear with no electron dense material content. Interestingly, exosome-like vesicles were never observed to be directly docked at the porosome base for fusion and release.

**Figure 2.**
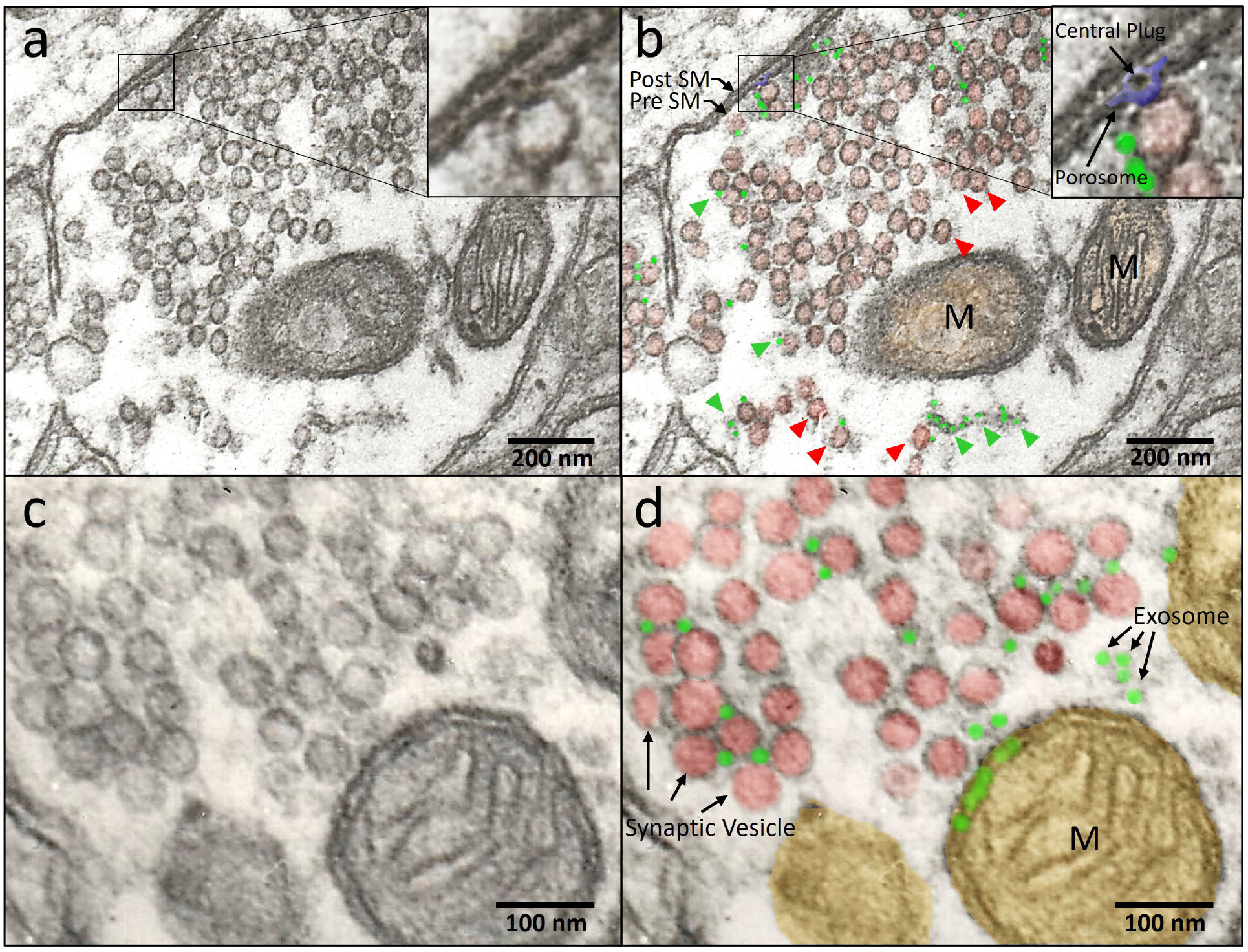
Electron micrograph (EM) of rat brain synaptosome showing the association of 15-20 nm exosome-like vesicles associated with synaptic vesicles, including docked synaptic vesicles at the neuronal porosome complex. Note exosome-like vesicles that appear to be released from the mitochondria present within the synaptosome compartment **(a-d).** Pseudo-color of the EM micrographs provide greater clarity.

### Electron microscopy on Min6 cells reveal the release of exosome-like vesicles via the porosome complex

In Min6 cells, multi-vesicular bodies are seen in electron micrographs. Occasionally, an insulin containing granule is found fused with a multi-vesicular body. Such a multi-vesicular body fused with an insulin containing granule, is seen docked and fused at the porosome base, releasing intact approximately 20 nm in diameter vesicles [Figure 3]. The porosome most likely is in the process of secreting insulin in addition to the release of exosome-like vesicles, since most of the electron dense material (insulin) from within the secretory granule has been released. These electron microscopy images further suggest that porosome complexes primarily involved in regulated secretion of hormones, enzymes and neurotransmitters, may also participate in the regulated release of exosomes and exosomal contents from cells. This hypothesis was further tested in Min6 cells.

**Figure 3.**
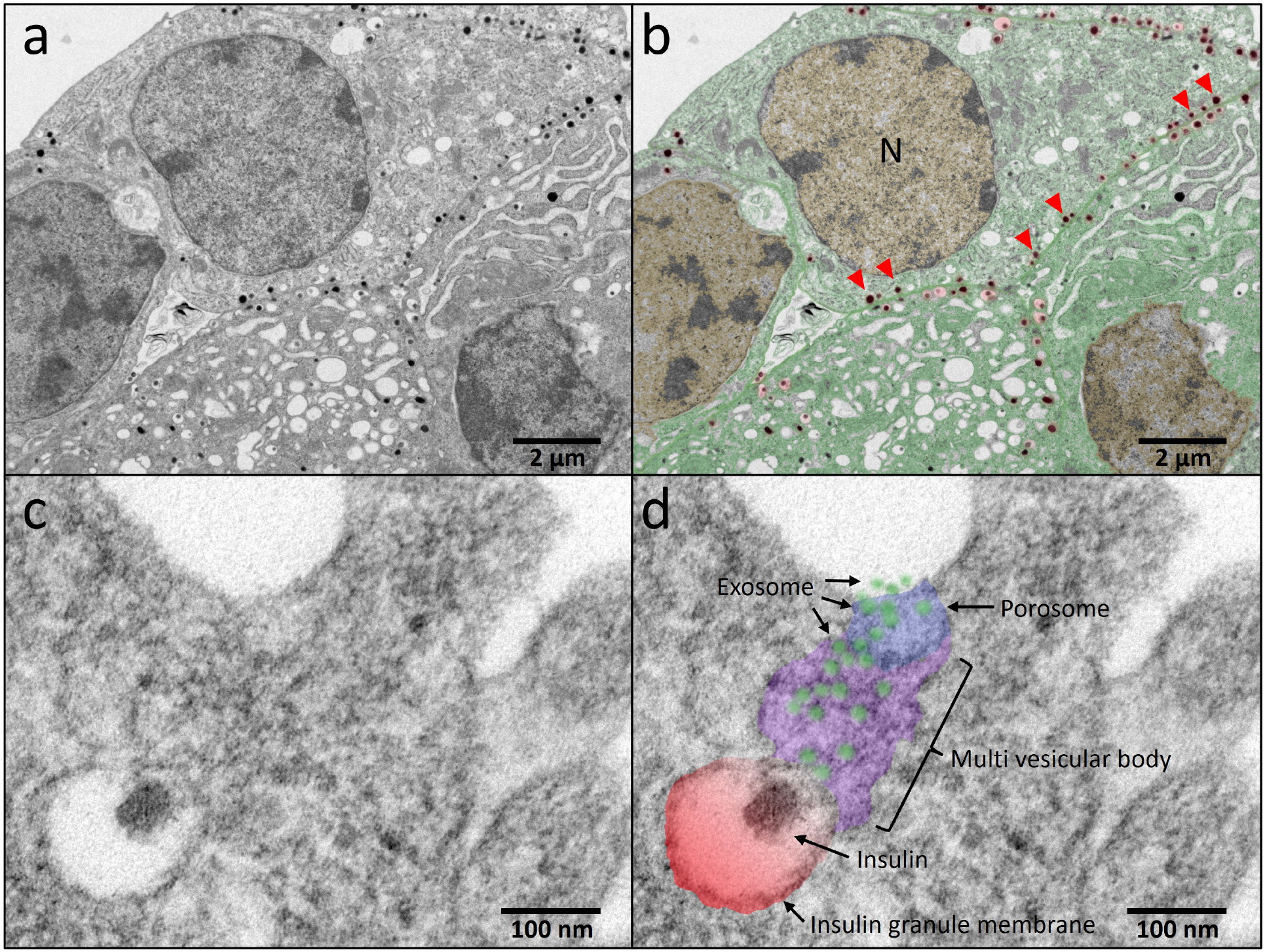
Electron micrograph showing release of exosome-like vesicles and the secretion of insulin via the porosome complex at the plasma membrane in mouse insulinoma Min6 cells. **(a, b)** Electron micrograph of a Min6 cell, and the nucleus (N), insulin containing electron dense secretory vesicles (red arrowhead) docked at the cell plasma membrane are observed. **(c, d)** A higher magnification electron micrograph demonstrates an insulin containing granule (red) fused with a multi-vesicular body (purple) containing exosomes (blue). The fused insulin granule and multi-vesicular body complex is seen docked at the porosome base and in the process of releasing exosome-like vesicles and insulin to the cell exterior. Note the 100 nm porosome complex, the 150 nm in diameter insulin granule, and the 15-20 nm exosomes being extruded from the cell via the porosome complex.

### Knockout of the porosome-associated protein gene ATP2C1 in Min6 cells result in loss of glucose-stimulated insulin secretion

CRISPR knockout (KO) of the porosome protein gene ATP2C1, results in the near absence of detectable levels of ATP2C1 gene product in the ATP2C1 KO Min6 cells [Figure 4]. The colocalization (yellow) of ATP2C1 (red, representing porosome) and insulin (green, representing insulin granules) in SCRM control cells, reflect the docked insulin granules at the porosome [Figure 4]. Very few such colocalizations are seen in the ATP2C1 KO Min6 cells. To further determine whether the expression and distribution of insulin and another porosome-associated protein Syntaxin 1 are affected, immunocytochemistry using Syntaxin 1- and insulin-specific antibody was performed [Figure 5]. Result from this study shows no detectable changes in either the expression or distribution and hence their colocalization of Syntaxin 1 and insulin.

**Figure 4.**
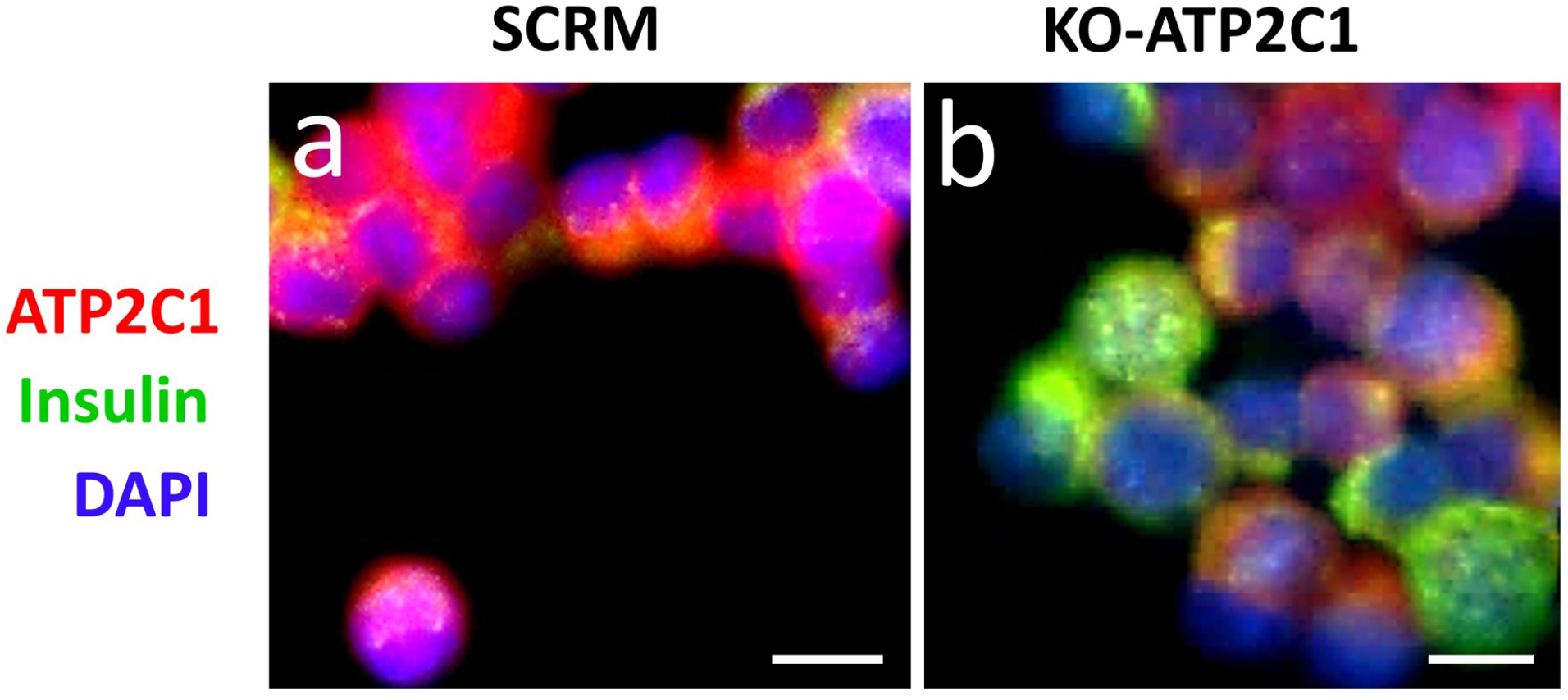
Immunofluorescent microscopy demonstrates the loss of ATP2C1 expression in ATP2C1 KO Min6 cells. Note however, that the expression and distribution of insulin and the porosome protein Syntaxin 1 in Min6 cells, or the morphology of the nucleus (DAPI blue) remain unaffected by the ATP2C1 KO. **(a)** Robust expression of ATP2C1 protein immunoreactivity is observed in the perinuclear Golgi region and at the cell periphery (plasma membrane) in SCRM control Min6 cells. Also note the yellow fluorescence, indicating co-localization of the porosome-associated ATP2C1 gene product and insulin containing granules in the control SCRM Min6 cells. In contrast in the ATP2C1 KO cells, the expression ATP2C1 is greatly reduced, resulting in an increase in undocked insulin containing granules (green) **(b)**. The DAPI stained nucleus and the overall cell morphology in both the SCRM and ATP2C1 KO appears indistinguishable. *Scale bar = 10 μm.*

**Figure 5.**
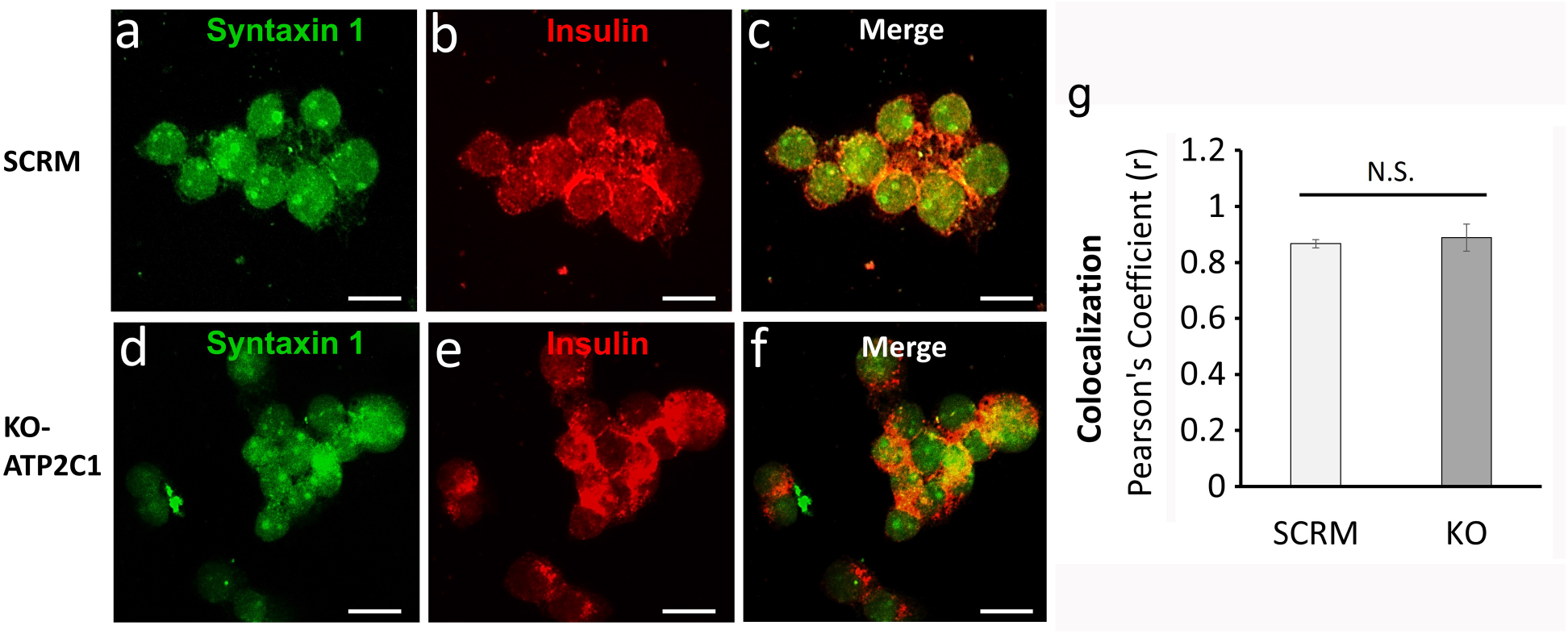
Immunofluorescent microscopy reflecting the unaffected expression and distribution of the porosome protein Syntaxin 1 and insulin containing secretory granules in SCRM control and in the ATP2C1 KO Min6 cells. **(a-f)** Note the circular patches of Syntaxin 1 immunoreactivity (porosomes) and the punctate insulin immunoreactivity representing the distribution of insulin containing granules. The merged images in (**c**) and (**d**) and their assessment using Pearson Correlation Coefficient **(g)**, reflect no difference between the SCRM controls and the ATP2C1 KO (n=3). These results further suggest that certain porosomes in Min6 cells may be devoid of the ATP2C1 gene product. *Scale bar = 20 μm.*

### Knockout of the porosome-associated protein gene ATP2C1 in Min6 cells, results in loss of glucose-stimulated insulin secretion and exosome release

Immunoblot analysis of the CRIPSR knockout (KO) of the porosome protein gene ATP2C1 in Min6 cells, demonstrates little expression of the ATP2C1 gene product [Figure 6a]. This further confirms the immunocytochemistry results reported in Figure 5. Exposure of cells to elevated glucose levels, demonstrate a time-dependent increase in insulin release in both the control and the experimental ATP2C1 KO Min6 cells [Figure 6b, c]. Although little change in the basal levels of insulin release is observed in Min6 cells (0 min), a significant (p<0.05) loss in glucose-stimulated insulin secretion is demonstrated after 30 min following exposure to elevated glucose over controls [Figure 6c]. Additionally, note a loss in the rate of glucose-stimulated insulin secretion from 0.08%/min in the SCRM control Min6 cells to 0.02%/min in the ATP2C1 KO Min6 cells. Secretion assays, besides demonstrating that ATP2C1 KO results in a loss of glucose-stimulated insulin secretion, also demonstrate a loss in exosome release [Figure 5b, c, d]. At the 30 min post glucose exposure, CD63 immunoreactivity is detected in Western blots on the resolved incubation medium from SCRM control Min6 cells, while in contrast, the ATP2C1 KO cells, exhibit no detectable signal [Figure 6d]. To test if cells are live and healthy, and to test if DNA is released during glucose stimulation of Min6 cells, the secretion assay medium from all three time points (0 min, 10 min and 30 min) in SCRM control and ATP2C1 KO was tested for DNA. Except for the standard (M), no detectable DNA bands are observed in the cell medium resolved using agarose gel followed by ethidium bromide staining for all time points in both the control and experimental Min6 cells [Figure 7]. These results demonstrate that the cells are live and healthy throughout the experiment, and that Min6 cells do not release DNA following a glucose challenge.

**Figure 6.**
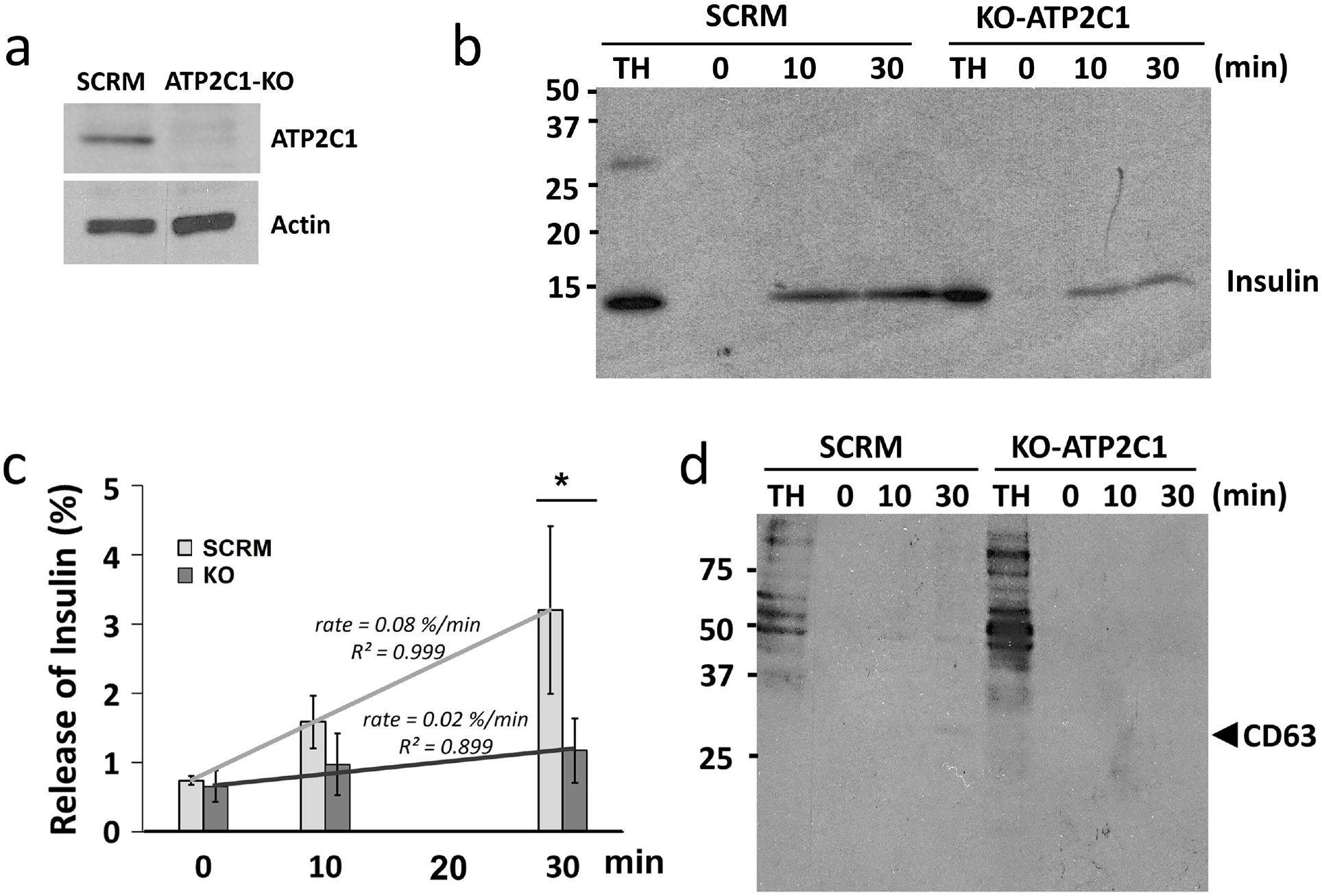
Immunoblot analysis demonstrates that there is significant loss of ATP2C1 expression in ATP2C1 KO Min6 cells, and that this absence of the porosome-associated protein results in a loss of both glucose-stimulated insulin secretion and exosome release. **(a)** Note the near absence of immunodetectable ATP2C1 gene product in the ATP2C1 KO Min6 cell homogenate compared to the SCRM control. **(b)** Immunoblot analysis of 5 μg of total Min6 cell homogenate (TH) from SCRM and ATP2C1 KO Min6 cells, and equal volumes of Min6 cell incubation medium (PBS) at different times following a glucose challenge, were resolved using SDS-PAGE followed by electrotransfer to nitrocellulose membrane and probed using the insulin-specific antibody and or the exosome-specific antibody CD63. Note the significant loss in both the potency and efficacy of glucose-stimulated insulin secretion in ATP2C1 KO Min6 cells compared to SCRM controls (n=5) **(b, c)**. Similarly, undetectable levels of the CD63 immunoreactivity in the ATP2C1 KO cell incubation media following 30 min of glucose stimulation is observed, which in contrast is clearly observed in the 30 min SCRM control medium (despite the high dilution).

**Figure 7.**
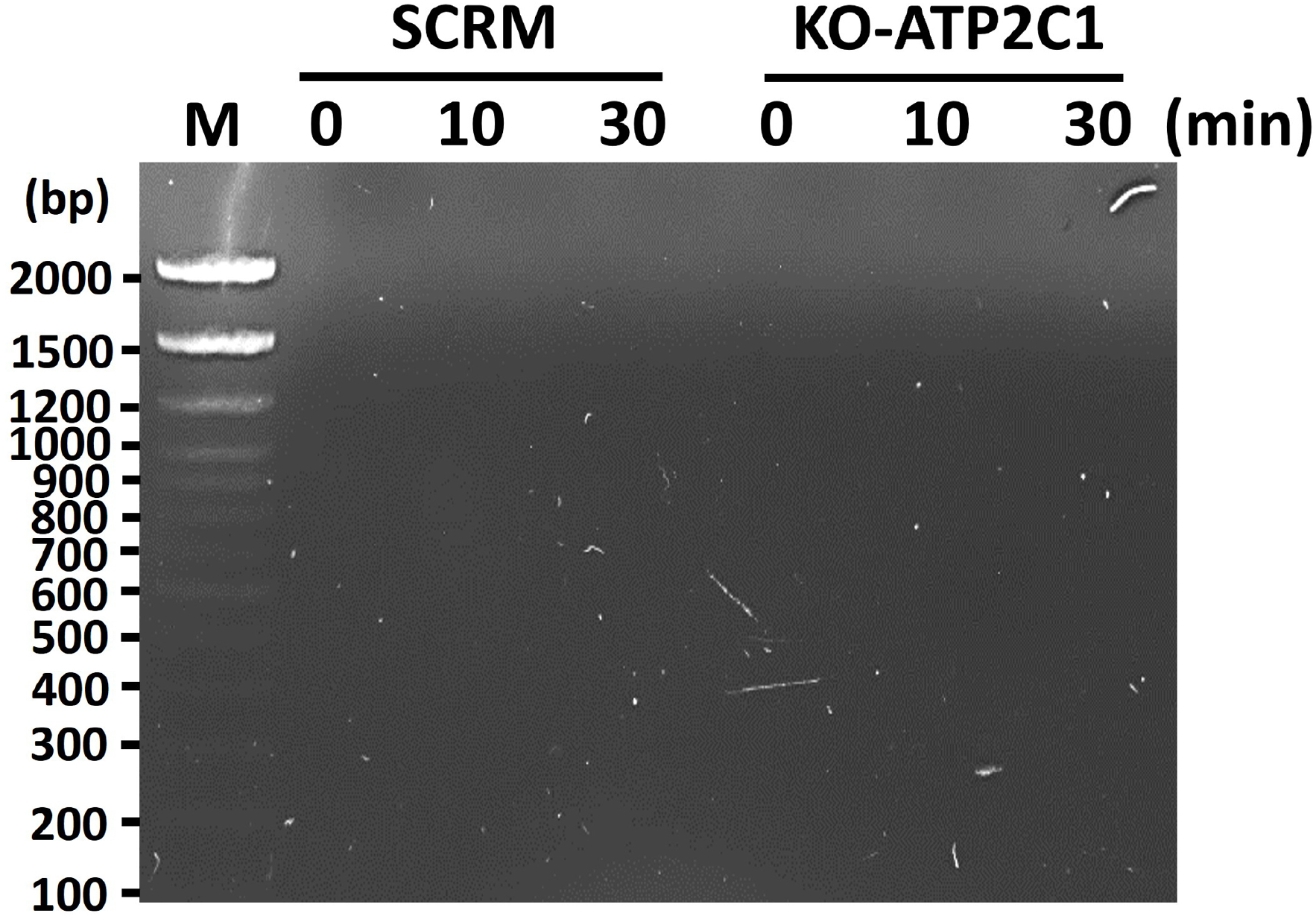
Min6 cells exposed to glucose for up to 30 min are healthy and intact, and do not release DNA. Agarose gel electrophoresis performed on a positive control DNA sample and Min6 cells (both SCRM and ATP2C1 KO) incubation medium following glucose challenge, demonstrate no release of the nucleotide at any time point post glucose challenge. These results additionally reflect that the viability and integrity of Min6 cells are intact throughout the course of the experiment.

Results from this study further establish the role of the porosome as a universal secretory portal in cells, that regulates the kiss-and-run mechanism of fractional insulin release, and further supports its role in the release of exosomes. Additional ongoing studies in the laboratory utilizing differential expansion microscopy (21) combined with immuno-electron microscopy, will provide additional clarity to this possible process of porosome-mediated exosome release and the chemistry and biogenesis of these 15-20 nm exosome-like vesicles.

## Acknowledgement

Work presented in this article was supported by the Viron Molecular Medicine Institute.

## Author Contributions

B.P.J. developed the idea, designed the experiments and wrote the paper. W-J.C. and B.F. performed the immunocytochemistry, CRISPR Knockout and Westerns; M.G.Z. performed electron microscopy on rat brain tissue; D.J.T performed electron microscopy on Min6 cells; I.G. participated in the idea development and image analysis. All authors participated in indepth discussions and proofreading the manuscript. The authors thank Daniel A. Walz for critical review of the manuscript.

## SUMMARY

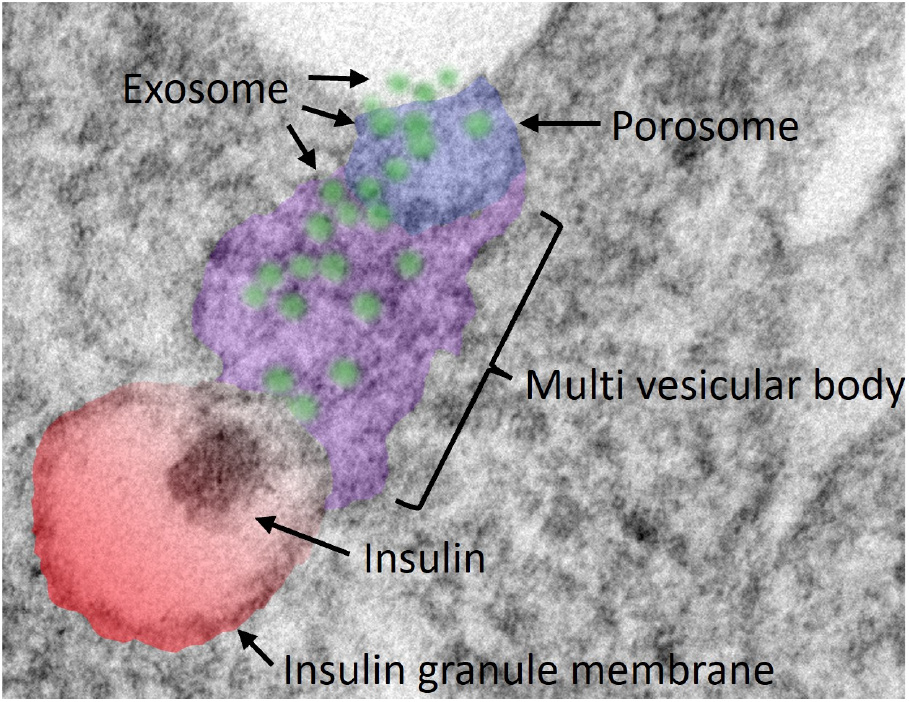

Exosomes and insulin are secreted through the porosome complex in mouse insulinoma Min6 cell. Electron micrograph demonstrates an insulin containing granule (red) fused with a multi-vesicular body (purple) containing exosomes (blue), docked at the porosome base and in the process of releasing exosomes to the cell exterior.

